# A comparison of two methods for estimating measurement repeatability in morphometric studies

**DOI:** 10.1101/2020.05.07.083428

**Authors:** Zachariah Wylde, Russell Bonduriansky

## Abstract

1. Measurement repeatability is often reported in morphometric studies as an index of the contribution of measurement error to trait measurements. However, the common method of remeasuring a mounted specimen fails to capture some components of measurement error, and could therefore yield inflated repeatability estimates. Remounting specimens between successive measurements is likely to provide more realistic estimates of repeatability, particularly for small structures that are difficult to measure.
2. Using measurements of 22 somatic and genitalic traits of the neriid fly *Telostylinus angusticollis*, we compared repeatability estimates obtained via remeasuring (where a mounted specimen is measured twice) versus remounting (where a specimen is remounted between measurements). We also asked whether the difference in repeatability estimates obtained via the two methods depends on trait size.
3. Repeatability estimates obtained via remounting were lower than estimates obtained via remeasuring for each of the 22 traits, and the difference between estimates obtained via the two methods was generally greater for small structures (genitalic traits) than for large structures (legs, wings).
4. Remounting specimens between successive measurements can provide more accurate estimates of measurement repeatability than remeasuring from a single mount, especially for small structures that are difficult to measure.

## Introduction

Repeatability is estimated from repeated measurements taken on several individuals, and is typically calculated as the intra-class correlation coefficient—i.e., the proportion of total variance that is attributable to individual identity (Lessells & Boag, 1987) (Sokal & Rohlf, 1995; Stoffel, Nakagawa, & Schielzeth, 2017). Repeatability is estimated for many different reasons (Wilson, 2018). In morphometric studies, repeatability is useful as a gauge of the contribution of measurement error to trait measurements, and therefore the statistical power of analyses involving those measurements (Bailey & Byrnes, 1990; Yezerinac, Lougheed, & Handford, 1992). In principle, if Trait A has greater measurement error than Trait B, Trait A will have lower repeatability than Trait B, and analyses of variation in Trait A will have lower statistical power. From a statistical standpoint, the calculation of repeatability has received considerable attention in the literature (Lessells & Boag 1987; (Altaye, Donner, & Klar, 2001; Shoukri & Donner, 2001; Ghosh & Das, 2003; Stoffel et al., 2017).

However, specimen handling and remeasurement methods can also affect repeatability estimates. For example, a study on the morphometrics of skeletal traits found that, as measuring technique improved through experience, measurement error declined (Yezerinac et al., 1992). The repeatability of trait measurements can depend on both intrinsic (e.g. genetic) and extrinsic (e.g. environmental) factors (Wilson, 2018). The way in which one handles and measures a specimen is likely to be an important, but somewhat overlooked extrinsic factor that also influences repeatability. Guidelines exist on the number of individuals and number of measurements per individual required to obtain precise estimates of repeatability (Wolak, Fairbairn, & Paulsen, 2012), but guidelines on appropriate specimen handling and remeasurement techniques are currently lacking.

To provide useful estimates of the contribution of measurement error to trait measurements, repeatability estimates must capture as many sources of measurement error as possible. In morphometric studies (particularly those involving small specimens, such as insect body parts), measurement error typically reflects how specimens are mounted, and how their dimensions are quantified. A common procedure is to remeasure a single mount (or an image of a single mount). This method has been used widely in studies by our lab and other groups (Blanckenhorn, Kraushaar, Teuschl, & Reim, 2004; Hosken, Minder, & Ward, 2005; Bertin & Fairbairn, 2007; Bonduriansky, 2007; Cayetano & Bonduriansky, 2015; Bonduriansky et al., 2015). Many other morphometric studies do not specify how repeatability was estimated, or do not report repeatability at all. However, this method omits sources of measurement error associated with specimen mounting. For example, each time a specimen is mounted its orientation relative to the focal plane of the microscope or camera will be slightly different, resulting in different degrees of parallax error. Soft specimens may also be distorted slightly each time they are handled, and images of specimens mounted in fluid medium (such as glycerol or saline solution) may be shadowed or distorted in various ways by the fluid.

Remounting specimens between successive measurement is therefore likely to yield better estimates of measurement repeatability. However, remounting takes time and effort, and it is not clear how substantially estimates of repeatability obtained by remounting would differ from estimates obtained by remeasuring the same mount, or whether the difference between estimates obtained by these methods varies between traits that can be measured with relatively little error (such as large, flat and stiff morphological structures) and traits that are subject to greater measurement error (such as small or soft morphological structures). To address these questions, we compared repeatability estimates obtained via remeasuring and remounting for 22 morphological traits of the neriid fly *Telostylinus angusticollis*. The traits included small and weakly sclerotized genitalic structures as well as relatively large, strongly sclerotized and flat structures (legs, wings).

## Material and methods

### STUDY SYSTEM

We utilised a morphometric dataset on the neriid fly *Telostylinus angusticollis* that included 22 somatic and genitalic traits measured on individuals reared on “rich” and “poor” larval diets (Wylde & Bonduriansky, submitted). Briefly, eggs collected from stock flies were reared using a nutrient-intermediate larval diet based on (Sentinella, Crean, & Bonduriansky, 2013), and randomly chosen adults were then paired to create 17 mating pairs. From each pair, 40 eggs were split equally between nutrient-poor and nutrient-rich larval diet treatments (Sentinella et al., 2013). Adults emerging from these larval diets were frozen for measurement ~24 h after emergence (i.e., once their exoskeletons had sclerotized fully). Larval diet manipulation influences adult body size and shape in *T. angusticollis* (Bonduriansky, 2007), and therefore increased the range of variation in the sizes of morphological traits examined in this study.

We measured six genitalic and 12 somatic traits on each of 93 males (n =43 poor diet, n = 50 rich diet), and four genitalic and 11 somatic traits on each of 96 females (n = 49 poor diet, n = 47 rich diet). All trait measurements were lengths in mm except for testis area, which was measured in mm^2^ (see Figure 1, 2 for definitions of trait measurements). Two methods were used to estimate repeatability for each trait. First, each specimen was mounted on the slide and imaged, and then remounted and reimaged; separate measurements were then made from the two images (’remounting’ method). Second, each trait was measured twice from a single image (‘remeasuring’ method). For the remeasuring method, we chose which of the two images to remeasure based on a random sequence of numbers generated from a binomial distribution.

**Fig. 1.**
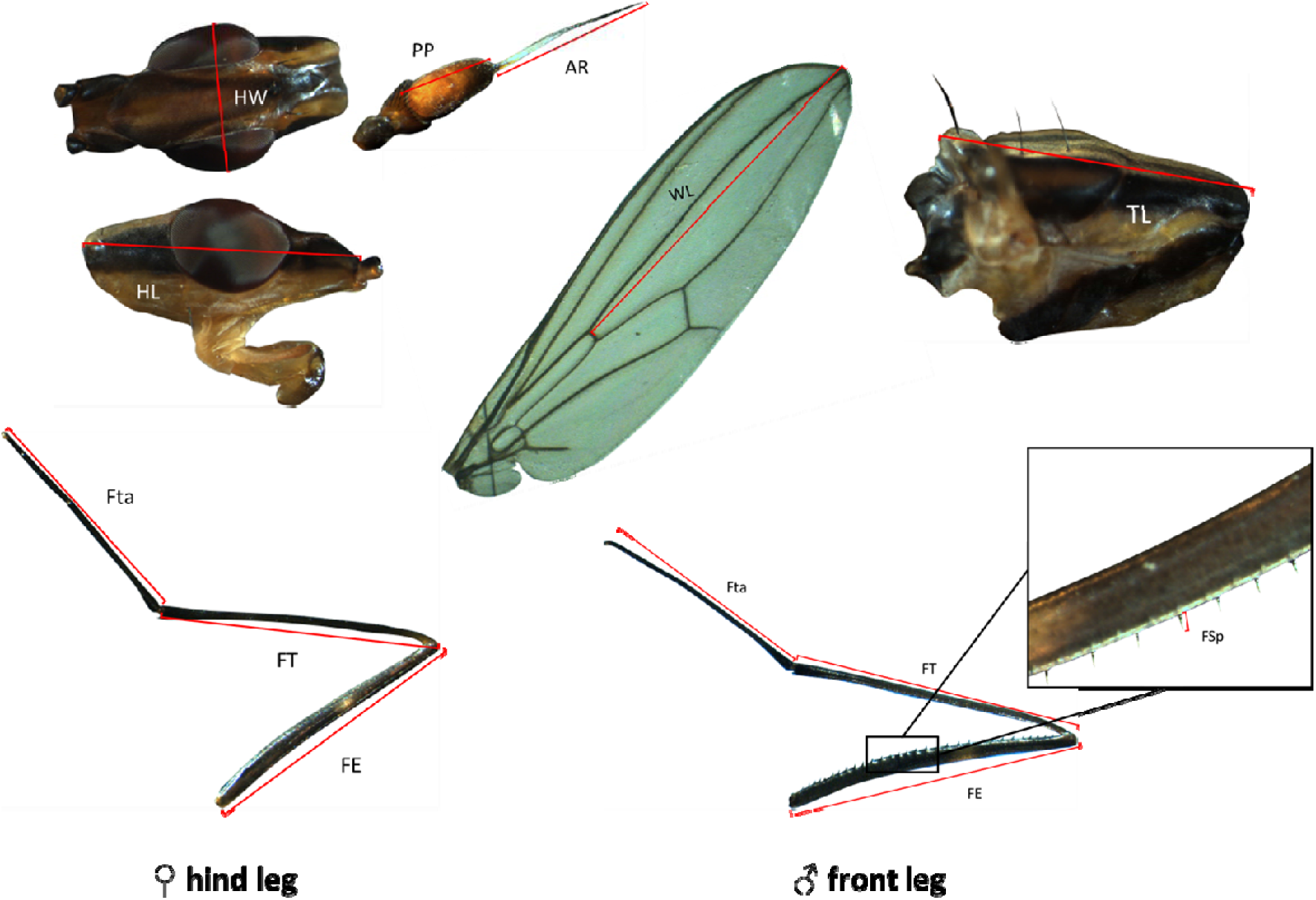
Larger sclerotized body parts of *T. angusticollis*. Shared traits between male and female include HW (head width), HL (head length), PP (post-pedicel length), AR (arista length), WL (Wing length) and TL (thorax length). In females the hind leg was measured, whereas in males the front leg was measured. Both legs were broken down into measurements of FE (femur length), FSp (femur spine length, males only), FT (tibia length), Fta (tarsus length). Trait images are not to scale.

**Fig. 2.**
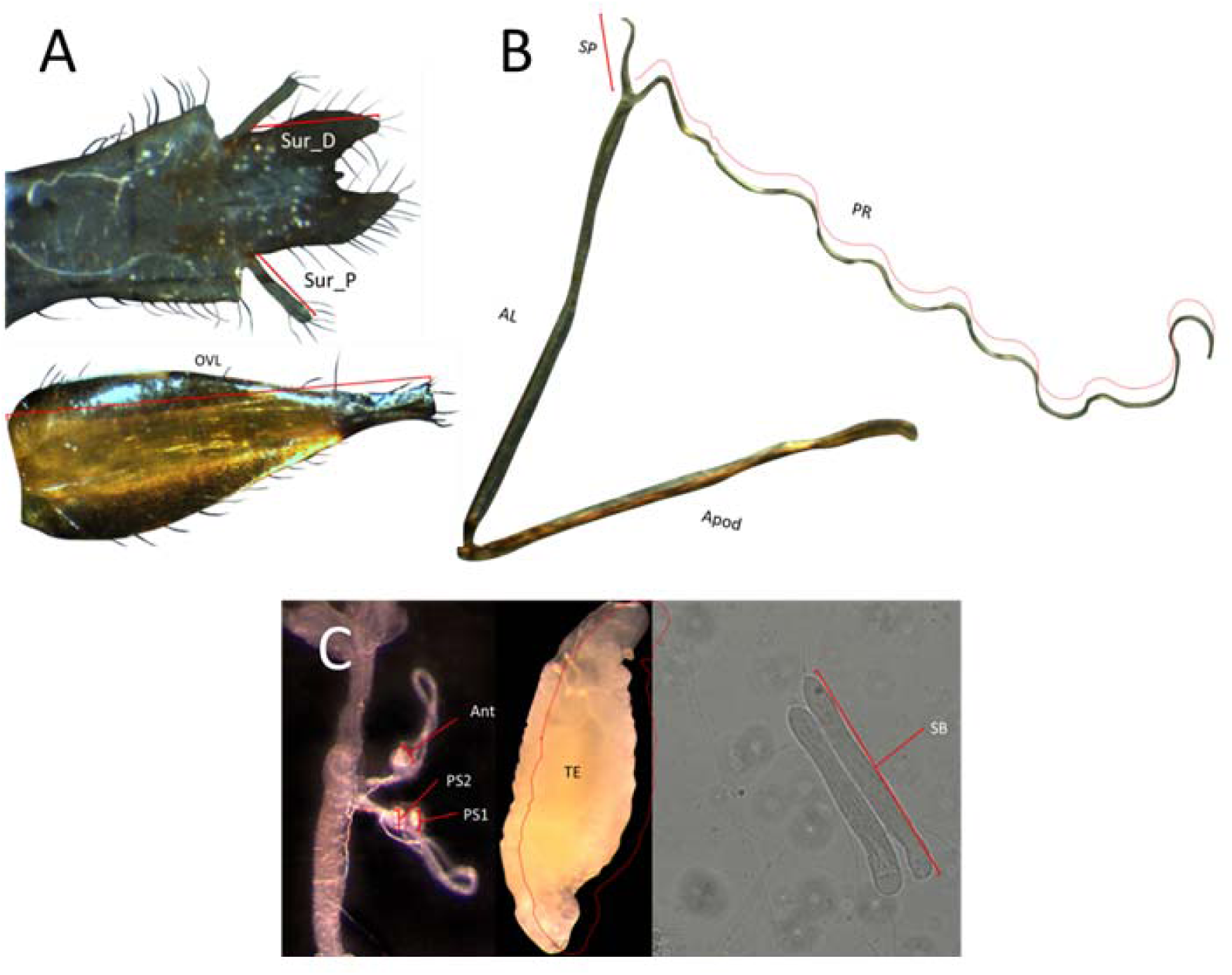
Genitalic and other smaller and unsclerotized traits of *T. angusticollis*. A: External genitalic traits: male Sur_P (proximal surstyli length) and Sur_D (distal surstyli length); female oviscape (OVL). B: Male genitalic apparatus: Apod (apodeme length), AL (Aedeagus length), SP (short anterior processus length), PR (processus length). C: left, female genitalic apparatus: PS1, PS2 (posterior spermathecae 1 & 2 width), Ant (anterior spermatheca width); male TE (testis, area mm^2^) and SB (sperm bundle length).

### SAMPLE PREPARATION

For each individual, the head, wings, legs and antennae were separated from the thorax and the genitalia were dissected out. Body parts were laid flat onto 1-1.2 mm microscope slides (ISSCO®) with an in-built micrometre for measurement calibration. To minimize parallax error, heads were positioned on slides covered with double-sided tape. Genitalic structures were mounted in 7.2 pH Phosphate Buffered Saline (PBS) and covered with 22 mm coverslips. The external genitalia (epandrium), male surstyli (proximal and distal) and internal section of the genitalia (carefully removed as one unit that included the apodeme, aedeagus and processes) were separated from the epandrium and placed under a coverslip. All somatic (both sexes) and male genitalic traits were imaged using a Leica MZ 16A stereoscope fitted with a Leica MC170 HD camera. Before dissection of spermathecae, the female oviscape was imaged and its length measured. The female reproductive tract with spermathecae was then carefully removed, cleaned and mounted in PBS as described above. Spermathecae were imaged using a Zeiss Axioskop 40 compound microscope fitted with a DinoEyepiece® camera at 20X magnification.

### STASTICAL ANALYSIS

All analyses were carried out using R 3.6.2 (R Core Team, 2019). Repeatability was calculated for all traits as the variance among individual trait means (individual-level variance VI) over the sum of individual-level and residual variance VR: R = VG/(VG + VR). We split the data by sex and method of measurement (remeasured or remounted) and fit separate linear mixed models using the packages “lmerTest” (Bates, Maechler, Bolker, & Walker, 2015; Kuznetsova, Brockhoff, & Christensen, 2017) and “lme4” (Bates et al., 2015) where trait size was the response variable, larval diet (rich or poor) was the fixed categorical predictor, and trait I.D. was the random effect. Subsequently, we used parametric bootstrapping (1000 iterations, 500 permutations) to obtain uncertainty in estimated repeatability values using the package “rptR” (Stoffel et al., 2017).

On the mean repeatability estimates for each trait, we conducted a paired Wilcoxon signed rank test to compare estimates obtained via the remounting vs. remeasuring methods. We then fit a linear mixed model to the mean repeatability estimates, with method (remeasuring vs. remounting) as a fixed categorical predictor, mean trait size as a fixed covariayte, and the two-way interaction between method and trait size and Trait_ID as the random factor. Conditional effect sizes reflect the variance explained by both the fixed and random effects (Nakagawa & Schielzeth, 2013) and allow us to quantify the magnitude of the influence that each factor has on the dependent variable (repeatability).Therefore, we also calculated the conditional effect sizes for each factor using three models that included either just method, mean trait size or the interaction between them using the technique outlined by Nakagawa & Schielzeth (2013).

## Results

Repeatability estimates obtained via remounting were lower than estimates obtained via remeasuring for each of the 22 traits (Table 1) (Wilcoxon signed-rank test, *w* = 496, *p* <0.001). The difference between repeatability estimates obtained via remeasuring and remounting methods increased as remounted repeatability decreased (*r* = −0.98, *t* = −28.1, *p*<0.001; Fig. 3). There was a significant interaction between trait size and measurement methodology whereby the difference between estimates obtained via remounting vs. remeasuring was greater for smaller traits (Table 2). As traits increased in size, and therefore decreased in average difficulty of consistent measurement, differences between the two methods decreased (Figure 4).

**Table 1.** Estimates of trait repeatability based on remounting and remeasuring methods for 22 morphometric traits of *Telostylinus angusticollis*. Traits are ordered by decreasing mean size (mm).

| Trait type | Trait | Sex | mean trait size (mm) | Repeatability (remeasured) | Repeatability (remounted) |
| --- | --- | --- | --- | --- | --- |
|  |  |  |  | [95% CI] | [95% CI] |
| Somatic | FSp | male | 0.051 | 0.919 [0.881, 0.946] | 0.777 [0.672, 0.859] |
| Genitalic | Ant | female | 0.063 | 0.995 [0.992, 0.997] | 0.901 [0.847, 0.939] |
| Genitalic | PS1 | female | 0.073 | 0.995 [0.993, 0.997] | 0.932 [0.892, 0.957] |
| Genitalic | PS2 | female | 0.075 | 0.994 [0.991, 0.996] | 0.919 [0.876, 0.949] |
| Somatic | SB | male | 0.161 | 0.994 [0.990, 0.996] | 0.760 [0.655, 0.847] |
| Genitalic | SP | male | 0.170 | 0.977 [0.964, 0.985] | 0.863 [0.794, 0.912] |
| Genitalic | Sur_P | male | 0.177 | 0.967 [0.949, 0.978] | 0.835 [0.750, 0.894] |
| Genitalic | Sur_D | male | 0.295 | 0.980 [0.970, 0.987] | 0.759 [0.642, 0.843] |
| Somatic | TE | male | 0.378 | 0.996 [0.995, 0.998] | 0.943 [0.906, 0.965] |
| Somatic | PP | female | 0.519 | 0.941 [0.914, 0.960] | 0.649 [0.501, 0.759] |
| Somatic | PP | male | 0.659 | 0.975 [0.963, 0.983] | 0.865 [0.795, 0.913] |
| Somatic | AR | female | 1.050 | 0.987 [0.980, 0.992] | 0.810 [0.728, 0.872] |
| Somatic | AR | male | 1.077 | 0.994 [0.991, 0.996] | 0.800 [0.690, 0.871] |
| Genitalic | Apod | male | 1.091 | 0.964 [0.946, 0.977] | 0.827 [0.736, 0.889] |
| Genitalic | AL | male | 1.136 | 0.996 [0.994, 0.998] | 0.808 [0.712, 0.879] |
| Somatic | HW | female | 1.309 | 0.993 [0.990, 0.996] | 0.861 [0.800, 0.906] |
| Somatic | HW | male | 1.316 | 0.993 [0.990, 0.996] | 0.865 [0.794, 0.912] |
| Somatic | HL | female | 1.705 | 0.993 [0.989, 0.995] | 0.854 [0.788, 0.903] |
| Genitalic | PR | male | 1.864 | 0.984 [0.976, 0.990] | 0.938 [0.902, 0.962] |
| Genitalic | OVL | female | 1.946 | 0.991 [0.987, 0.994] | 0.929 [0.893, 0.952] |
| Somatic | HL | male | 2.240 | 0.998 [0.997, 0.999] | 0.922 [0.881, 0.950] |
| Somatic | TL | female | 2.340 | 0.994 [0.991, 0.996] | 0.927 [0.889, 0.953] |
| Somatic | TL | male | 2.545 | 0.997 [0.996, 0.998] | 0.959 [0.937, 0.974] |
| Somatic | Fta | Female | 2.589 | 0.992 [0.988, 0.995] | 0.968 [0.950, 0.980] |
| Somatic | WL | Female | 3.460 | 0.999 [0.999, 0.999] | 0.990 [0.984, 0.994] |
| Somatic | Fta | male | 3.583 | 0.993 [0.989, 0.996] | 0.953 [0.925, 0.972] |
| Somatic | WL | male | 3.671 | 0.998 [0.997, 0.999] | 0.977 [0.964, 0.986] |
| Somatic | FT | female | 3.743 | 0.992 [0.988, 0.995] | 0.959 [0.939, 0.973] |
| Somatic | FE | female | 4.354 | 0.997 [0.996, 0.998] | 0.976 [0.964, 0.985] |
| Somatic | FT | male | 4.877 | 0.988 [0.982, 0.992] | 0.961 [0.938, 0.975] |
| Somatic | FE | male | 4.946 | 0.999 [0.998, 0.999] | 0.982 [0.972, 0.989] |

**Table 2.** Generalized linear mixed effects model of repeatability with conditional effect sizes.

|  | Estimate | Std. Error | *df* | *F* | *p* | R<sup>2</sup>GLMM(*c*) |
| --- | --- | --- | --- | --- | --- | --- |
| (Intercept) | 0.980 | 0.013 | 46.146 | - | - | - |
| Mean trait size | -0.155 | 0.016 | 36.412 | 95.578 | <0.001 | 15.84 % |
| Method | 0.004 | 0.006 | 36.528 | 18.147 | 0.497 | 59.84% |
| Method: Mean trait size | 0.032 | 0.007 | 36.412 | 21.039 | <0.001 | 72.66% |

**Fig. 3.**
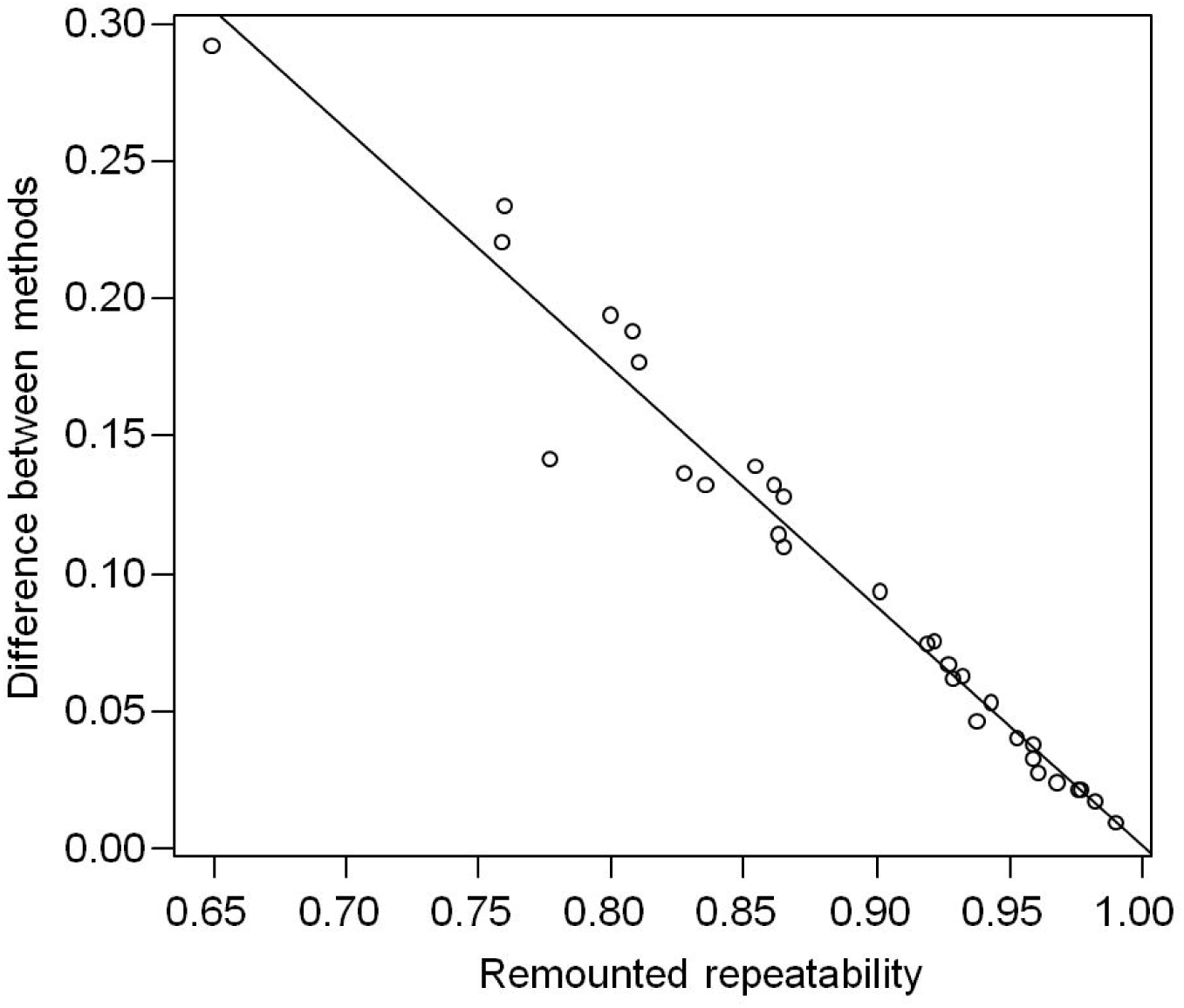
The difference between repeatability estimates obtained via remeasuring and remounting methods increases as remounted repeatability decreases.

**Fig. 4.**
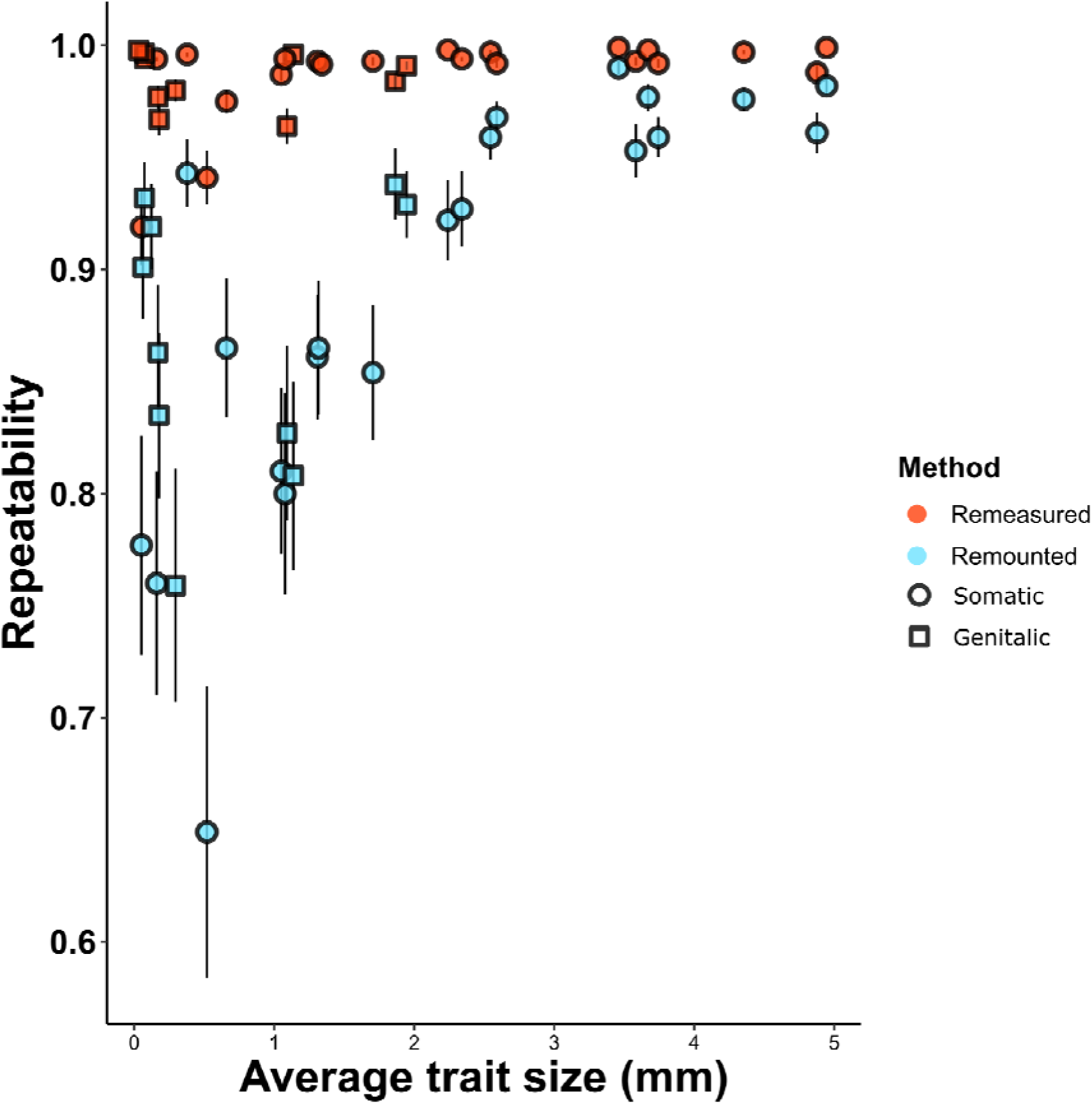
Estimates of repeatability obtained by remounting and remeasuring methods, ranked by mean trait size (smallest to largest, left to right). Bars represent standard error of the mean calculated from bootstrapped estimates of repeatability.

## Discussion

It is clear from our results that both the sample-handling method and the nature of the trait itself can strongly influence estimates of repeatability. For each of the 22 traits examined, we found that repeatability estimates obtained by remounting samples between successive measurements were smaller than repeatability estimates obtained by remeasuring a single mount. Moreover, the difference between the estimates obtained via the two methods increased on average as trait size decreased. Small, soft, or rounded body parts are likely to be subject to greater measurement error than large, hard or flat parts, and remeasuring a single mount is likely to substantially underestimate measurement error for such traits and yield substantially inflated estimates of measurement repeatability. The inflation of repeatability can have important consequences, potentially leading to overestimates of statistical power or parameters of interest such as heritability.

Repeatability is often used to quantify the accuracy and consistency of phenotypic measurements in evolutionary and behavioural ecology. In morphometric studies, repeatabilities are often reported as a guide to the signal-to-noise ratio of various trait measurements. This can be useful in gauging whether a stronger effect for a particular trait, relative to other traits, might simply reflect differences among traits in measurement error e.g. (Cassidy, Bath, Chenoweth, & Bonduriansky, 2013). Repeatability estimates are also sometimes used to estimate the upper bounds of narrow- and broad-sense heritability (Boake, 1989; Falconer & Mackay, 1996; Lynch & Walsh, 1998; Dohm, 2002). Furthermore, repeatabilities have been used to test predictions of condition-dependent models of sexual selection. For example, Foley et al (2012) found that for cervid antler traits, repeatability declines as environmental variation increases, supporting the idea that antlers serve as an honest signal of individual condition. Our findings suggest that estimates of repeatability obtained by remounting specimens between successive measurements are more accurate, and that the importance of using this method, rather than simply remeasuring a single mount or image, increases as trait measurement error increases. Interestingly, while the difference between methods was small for all of the large structures in our sample (i.e., structures > 2.5 mm in length), the difference between methods varied considerably for the smaller structures (Fig. 4). This suggests that trait size does not account fully for variation in measurement error and that other factors (e.g. shape or degree of sclerotization) could also be important.

Although we have focused in this study on morphometric repeatability, the principles that we discuss generalize to research on other types of traits for which repeatability is quantified. In all studies that report repeatability, it is important to consider the major sources of measurement error or variability, and how to estimate repeatability in a way that captures these factors (Wilson 2018). For example, many studies estimate repeatability for individual behaviour as a way to describe ‘behavioural syndromes’ or ‘animal personalities’ (Sih, Bell, & Johnson, 2004; Bell, Hankison, & Laskowski, 2009). In such studies, rather than retesting individual responses under identical conditions, it may be more relevant to test responses across different types of arenas or social environments. Behaviours remeasured in different contexts will probably be less repeatable (Fisher, James, Rodríguez-Muñoz, & Tregenza, 2015; Arvidsson, Adriaensen, van Dongen, De Stobbeleere, & Matthysen, 2017), but such estimates may be more meaningful if the aim is to quantify the consistency in individual responses across the range of contexts where the behaviour is expressed. For example, if the aim is to estimate the repeatability of a behavioural courtship signal, then it may be useful to measure this signal in multiple contexts where this signal is employed. Simply remeasuring the signal within the same context may yield a misleading estimate of signal repeatability.

In conclusion, our findings suggest that remeasuring from the same mount can yield strongly inflated repeatability estimates in morphometric studies, especially for traits subject to large measurement error. Remounting samples between measurements is likely to provide more meaningful estimates of repeatability. More broadly, the methods used to estimate repeatability should capture as many important sources of measurement error or variability as possible.

## Data accessibility

All data will be available on the Dryad data repository.

